# Multi-modal imaging of a single mouse brain over five orders of magnitude of resolution

**DOI:** 10.1101/2020.10.07.329789

**Authors:** Sean Foxley, Vandana Sampathkumar, Vincent De Andrade, Scott Trinkle, Anastasia Sorokina, Katrina Norwood, Patrick La Riviere, Narayanan Kasthuri

## Abstract

Mammalian neurons operate at length scales spanning five orders of magnitude; micron-scale-diameter myelinated axons project millimeters across brain regions, ultimately forming nanometer scale synapses on individual post-synaptic neurons. Capturing these anatomical features across that breadth of scale has required imaging samples with multiple independent imaging modalities (e.g. MRI, electron microscopy, etc.). Translating between the different modalities, however, requires imaging the *same* brain with each. Here, we imaged the same postmortem mouse brain over five orders of spatial resolution using MRI, whole brain micron-scale synchrotron x-ray tomography (μCT), and large volume automated serial electron microscopy. Using this pipeline, we can track individual myelinated axons previously relegated to axon bundles in diffusion tensor MRI or arbitrarily trace neurons and their processes brain-wide and identify individual synapses on them. This pipeline provides both an unprecedented look across a single brain’s multi-scaled organization as well as a vehicle for studying the brain’s multi-scale pathologies.

## 1. INTRODUCTION

The architecture of the mammalian brain is complicated, with brains operating on multiple scales spanning orders of spatial magnitude (Lichtman and Denk, 2011). Neurons in the brain project macroscopic distances (millimeters to centimeters) (S.R., 1899), while simultaneously producing microscopic connections and plasticity-related changes at the nanoscale (Grutzendler et al., 2002).

No current single existing imaging modality spans the six orders of length-scale (mm to nm) required to map the whole brain while simultaneously resolving individual synapses. Magnetic resonance imaging (MRI) allows *in vivo* (Basser et al., 2000) and postmortem (Dyrby et al., 2007; Miller et al., 2011) mapping of neuronal tracts spanning macroscopic distances but cannot easily observe individual neurons much less identify individual neuronal connections. At the other end of the resolution scale, advances in automated electron microscopy (EM) provide synapse-level resolution (3 nm) (Helmstaedter, 2013) but remain limited by serious computational challenges; a single voxel of MRI data (~1mm^3^) is almost 2 million terabytes of raw EM image data (Bouchard et al., 2016). Thus, electron microscopy remains best suited for neuron ‘connectomics’ - reconstructing connections of individual neurons over limited ranges of 100s of microns. A pipeline bridging these approaches would have to solve the problems of ‘context’; providing sufficient information and volume to help register an EM dataset in the context of an MRI volume. An ideal intermediate imaging modality would (a) achieve micron-scale resolution throughout entire an large volume tissue sample and (b) use a sample preparation compatible with a pipeline including MRI and electron microscopy.

Synchrotron-based x-ray tomography (μCT) nondestructively provides. micron-scale resolution over ~1cm^3^ whole tissue volumes - the size of a small mammalian brain - bridging the resolution disparity between MRI and EM imaging techniques (Fig. 1). Sample preparation for μCT uses aldehyde fixed material, compatible with postmortem MRI (Shepherd et al., 2009), followed by osmium staining and plastic embedding, compatible with large volume electron microscopy (Dyer et al., 2017). Thus, MRI imaging followed by μCT and EM provides a viable pipeline for multi-scale whole brain imaging. In this work we demonstrate the feasibility of such a pipeline by imaging a fixed postmortem mouse brain with MRI, preparing the whole brain for subsequent μCT and EM microscopy (Mikula et al., 2012; Mikula and Denk, 2015), re-imaging the entire brain with micron-resolution μCT (Dyer et al., 2017; Vescovi et al., 2018), and, finally, imaging isolated sub-volumes with automated serial EM (Kasthuri et al., 2015).

**FIGURE 1.**
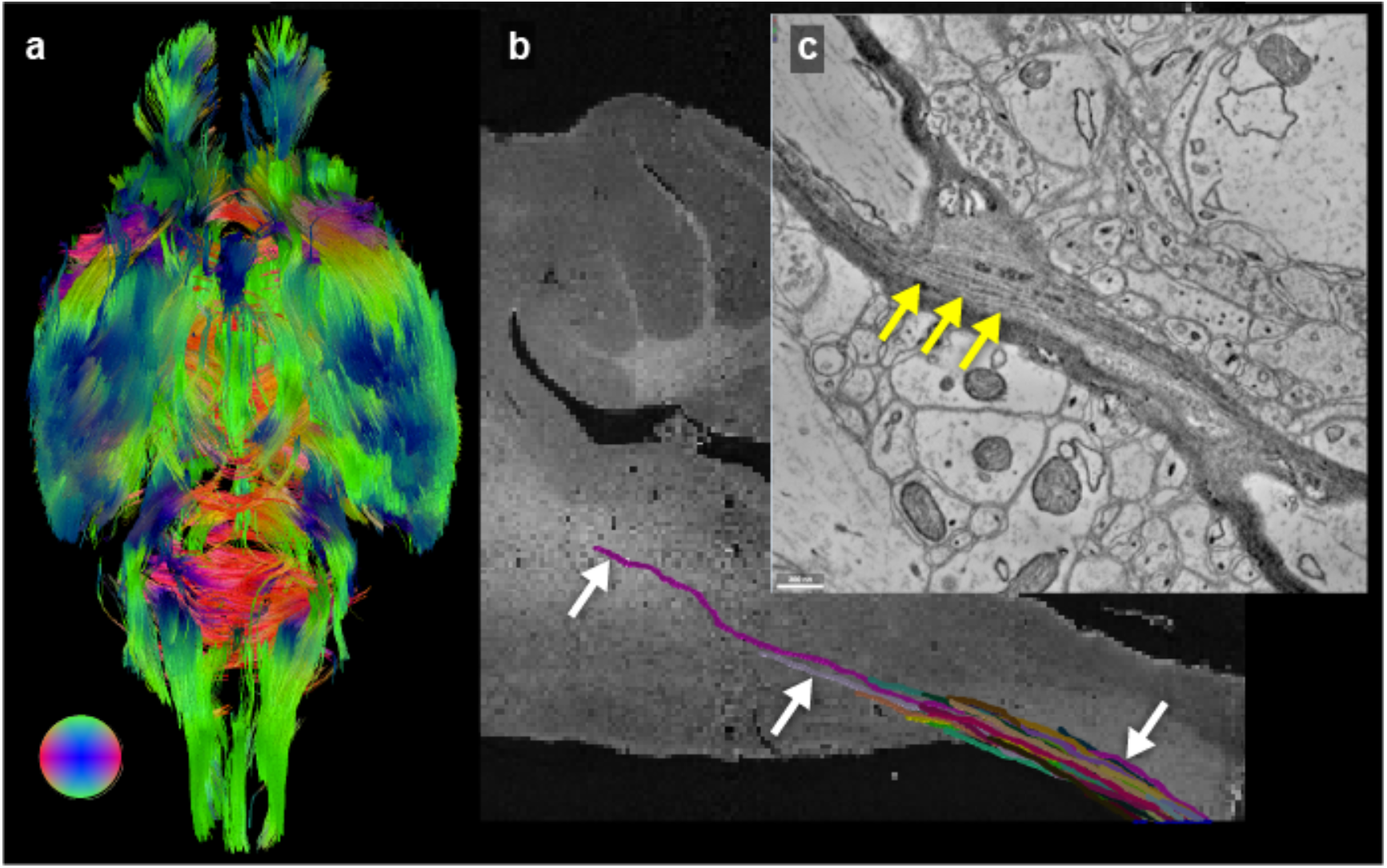
Imaging modality scale differences capture scale dependent information. By using an imaging pipeline of MRI, μCT, and EM, we can simultaneously resolve brain structures, like the white matter, at macro-, meso-, and microscopic-scales in the same brain. (a) We visualized whole brain interconnectivity by inferring white matter tracts from MRI diffusion tensor imaging. The colors denote tract directions - see inset color sphere. (b) We can trace individual axons (white arrows) within the same myelinated tracts over long distances with μCT, directly comparing axon paths with tracts produced in MRI. With the addition of EM, we can re-inspect the ultra-structure of individually traced axons from (b), such as (c) the individual wraps of the myelin sheath (yellow arrows).

## 2. MATERIALS AND METHODS

### 2.1 Sample preparation

All procedures performed on animals followed protocols approved by the Institutional Animal Care and Use Committee and were in compliance with the Animal Welfare Act and the NIH Guide for the Care and Use of Laboratory Animals. An eight week old, female C57bl6 mouse was deeply anesthetized with 60 mg/kg pentobarbital. The mouse was transcardially perfused with a solution (pH 7.4) of 0.1M Sodium Cacodylate and heparin (15 units/ml) immediately followed by a solution of 2% paraformaldehyde, 2.5% glutaraldehyde, and 0.1M Sodium Cacodylate (pH 7.4). The brain was carefully removed from the skull and post-fixed in the same fixative for 48h at 4°C. The order of serial imaging was magnetic resonance imaging (MRI), synchrotron-based x-ray tomography (μCT), and then electron microscopy (EM).

#### 2.1.1 MRI

The resected and aldehyde fixed brain was washed in PBS for 72h to remove fixative, which can be deleterious to MRI imaging (Dyrby et al., 2007; Shepherd et al., 2009). Just prior to MR imaging, brains were dried of excess PBS and placed in 10 ml falcon tubes. Tubes were filled with Fluorinert (FC-3283, 3M Electronics) for susceptibility matching and to improve shimming.

#### 2.1.2 μCT and Electron Microscopy

After MRI imaging the whole brain was transferred to 0.1M cacodylate buffer. Whole brain staining with osmium and other heavy metals and whole brain dehydration and plastic embedding were done as described (Mikula et al., 2012; Mikula and Denk, 2015). The brain was stained with multiple rounds of osmium and reduced osmium followed by dehydration and plastic embedding (Mikula and Denk, 2015).

### 2.2 Acquisition protocols

#### 2.2.1 MRI

Data were acquired at 9.4 Tesla (20 cm internal diameter, horizontal bore, Bruker BioSpec Small Animal MR System, Bruker Biospin, Billerica, MA) using a 6cm high performance gradient insert (maximum gradient strength: 1000 mT/m, Bruker Biospin) and a 35mm internal diameter quadrature volume coil (Rapid MR International, Columbus, Ohio). The brain was aligned such that the anterior/posterior portion of the olfactory limb of the anterior commissure (AC) was approximately parallel to the magnetic field of the scanner and the hemispheric midline was parallel to the scanner YZ plane.

Third order shimming was iteratively performed over an ellipse that encompassed the entire brain, but did not extend beyond the boundaries of the falcon tube/Fluorinert interface, using the Paravision mapshim protocol. B0 maps were produced by recording the voxel-wise frequency of the peak of the resonance, including additional sub-spectral resolution frequency produced by estimating the maximum peak amplitude of the resonance, described below. This was consistent with previously reported work (Foxley et al., 2015; Foxley et al., 2018), which described a high degree of field homogeneity across samples.

DTI was performed using a standard diffusion-weighted 3D spin echo sequence (TR = 400ms, TE = 18.5ms, b value = 3000 s/mm^2^, d = 5ms, D = 11.04ms, spatial resolution = 150um isotropic, number of b0’s = 16, number of non-collinear directions = 30, receiver bandwidth = 200kHz, partial Fourier along first phase encoding direction = 7/8, duration = 55hrs 19min 40sec).

Structural MRI data were acquired using a multi-echo gradient echo sequence. This produces a 4D dataset, where data are sampled along time in each spatial voxel to acquire the free induction decay (FID). Sequence parameters were chosen so that the entire voxel-wise FID was sampled to the noise floor (TR = 1000 ms, TE of first echo = 2.74ms, echo spacing = 2.74ms, number of echoes = 192, receiver bandwidth = 75kHz, flip angle = 68°, 100mm isotropic resolution, 4 averages, duration = 12 hours).

#### 2.2.2 μCT

The microCT data were acquired at the 32-ID beamline at the Advanced Photon Source, Argonne National Laboratory. The setup consists of a 1.8 cm-period undulator operated at a low deflection parameter value of *K* = 0.26. This yields a single quasi-monochomatic peak of energy 25 keV without the losses incurred by use of a crystal monochromator. For a sample 68 m from the undulator, this produces a photon fluence rate of about 1.8 x 10^7^ photons s^-1^ μm^-2^.

The x-rays were imaged using a 10 μm thick thin-film LuAG:Ce scintillator producing visible-light images then magnified using a 5X long Mitutoyo long working distance microscope objective onto a 1920 × 1200 pixel CMOS camera (Point Gray GS3-U3-51S5M-C). The effective object space pixel size is 1.17 μm and the field of view 2.25 × 1.41 mm^2^. The thickness of the thin-film scintillator matches the depth of focus of the objective lens, achieving a spatial resolution equivalent to the resolving power of the lens (1.3 μm for a NA of 0.21). Since the camera field of view (FOV) is substantially smaller than the mouse brain, we employed a mosaic strategy (Vescovi et al., 2018).

The sample was mounted on an air-bearing rotary stage (PI-Micos UPR-160 AIR) with motorized *x/y* translation stages located underneath and *x/y* piezo stages on top. Typical exposure times for a single projection image at one mosaic grid point and one rotation angle were 30 ms, and 3600 rotation angles were used at each grid point. The sample was translated through a 6 × the precomputed source link18 tomosaic grid.

#### 2.2.3 Electron microscopy

Regions-of-interest (ROI) in the brainstem and cerebellum were identified in the μCT data. The sample block was trimmed to each ROI and 50nm thick serial sections were collected on aluminium tape, attached to a wafer (Kasthuri et al., 2015), and imaged in an Apreo scanning electron microscope (SEM) (ThermoFisher Scientific). Images were obtained with 135nm or 3 nm in-plane resolution with 3μs dwell times.

### 2.3 Data processing and analysis

#### 2.3.1 MRI

Data were reconstructed, processed, and analyzed with IDL 8.2 (ITT Visual Information Solutions, Boulder CO), Matlab 2014b (The MathWorks Inc., Natick, MA, 2012), and FSL 5.0.9 (FMRIB Software Library, FMRIB, Oxford, UK).

3D multiple-gradient echo data were reconstructed to produce voxel-wise water spectra. Each complex 4D dataset (*x* × *y* × *z* × *t*) was Fourier transformed along time to produce three spatial dimensions and one spectral dimension (*x* × *y* × *z* × *t*). A T2*-weighted structural image was constructed from the spectral data, where image contrast was produced by isolating the maximum voxel-wise signal amplitude of the water spectrum (Al-Hallaq et al., 2002). This was achieved by identifying and storing the voxel-wise maximum peak signal amplitude in a 3D array with the same spatial dimensions as the acquisition. The mean b = 0 s/mm^2^ dataset from the diffusion acquisition was registered to the structural image via affine transformation using the FSL linear registration tool (FLIRT) (Jenkinson and Smith, 2001). Subsequent diffusion data were identically registered using the resultant transformation matrix. Principal, secondary, and tertiary diffusion directions were estimated using BEDPOSTX (Behrens et al., 2003).

#### 2.3.2 μCT

The acquired sub-sinograms were registered with sub-pixel shifting and stitched through pyramid blending into a complete sinogram. The center of rotation of the composite sinogram was estimated using an entropy-based approach. Reconstruction of the entire dataset in whole-block mode is performed using the gridrec implementation of direct Fourier reconstruction implemented in the TomoPy package resulting in a 11.7 × 11.7 × 17.7 mm^3^ reconstructed image volume at 1.17 μm isotropic voxels. The reconstructed volume was trimmed and stored in 8-bit format.

Following reconstruction, μCT data were visualized using Neuroglancer (https://github.com/google/neuroglancer), a high-performance WebGL based viewer for large volumetric datasets. Neuroglancer allows for memory-efficient loading of cross-sectional displays across arbitrary planes in the data. This allowed for simple, fine-tuned manual localization of the visible structures in the EM sections within the μCT dataset, using large blood vessels and neuronal somas as landmarks. Our μCT data can be publicly viewed online through Neuroglancer (http://neuroglancer-demo.appspot.com/) using http://nova.kasthurilab.com:8000/neuroglancer/recon_crop8_neurog/image as the precomputed source link.

All manually labeling of somas and myelinated axons was performed using Knossos (Helmstaedter et al., 2011).

## 3. RESULTS

### 3.1 Whole brain imaging with μCT

Previous work has shown the compatibility of MRI and EM in postmortem tissue samples (Liu et al., 2011). In order to assess the feasibility of utilizing μCT as a compatible bridge between the two, we developed a protocol for imaging entire mouse brains (1 cm^3^) at micron resolution with μCT. We leveraged advances in osmium staining and embedding an entire mouse brain (Mikula et al., 2012; Mikula and Denk, 2015), tomographic imaging extending the field of view of an X-ray imaging system (Vescovi et al., 2018), and parallelization of x-ray reconstruction algorithms on national lab supercomputers (Argonne National Laboratory) for creating contiguous image stacks from the ~10 terabyte sized raw ‘sinogram’ data. We were able to quickly image (~8 hours) and reconstruct an entire mouse brain at 1-micron resolution (Fig. 2a). The resulting series of images formed a well-aligned isotropic dataset with sufficient resolution and contrast to capture both individual anatomic regions (Fig. 2b; the full dataset can be explored in Supplemental Movie 1) as well as underlying brain wide cytoarchitecture. For example, both neuronal somas (Fig. 2c) and long distance myelinated axons bundle (Fig. 2d) could be clearly visualized in the same brain.

**FIGURE 2.**
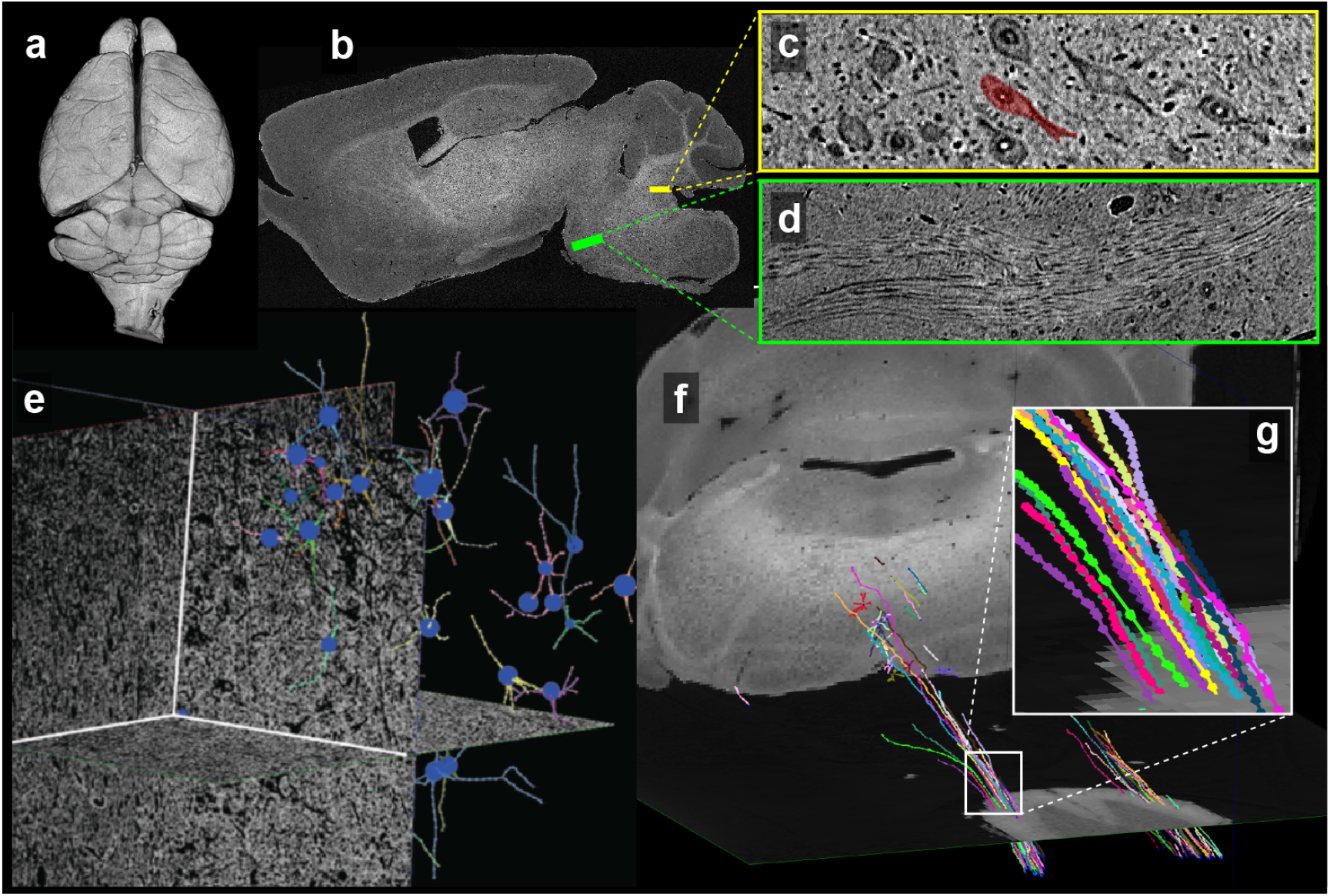
An entire mouse brain imaged at 1 micron resolution with μCT. We used high energy X-rays (23 kEV) for tomographic imaging of (a) a whole, intact mouse brain at ~1micron resolution, collecting ~10 terabytes of data in ~7 hours. A single downsampled sagittal slice (b), reveals obvious brain regions and myelinated tracts while the full resolution data contains enough detail and contrast to identify (c) individual neuronal soma, nucleus and nucleolus as well as (d) individual myelinated axons. Since μCT data are collected non-destructively, over large volumes, and with isotropic resolution, we can trace somas and their neuronal processes (dendritic trees, e) and individual myelinated axons (g) over long distances (f).

Data produced sufficient detail and contrast to trace cells, their morphologies, and individual myelinated axons several millimeters (Fig. 2e-g; Supplemental Movie 2; a down-sampled version can be viewed here: https://tinyurl.com/yd9c4jdq.).

### 3.2 Imaging a single sample through the complete pipeline

As described in the Methods section, a perfusion fixed rat brain was imaged with

MRI, prepared for microscopy, and imaged with both μCT and automatic tape-collecting ultra-microtome (ATUM) serial EM (Kasthuri et al., 2015). Thus, resolutions from each modality spanned 5 orders of magnitude on the same brain: 150μm DTI and 50μm structural MRI data, 1.2 μm μCT data, and 3 nm EM data. Typical results (Fig. 3) show similar regions from the same brain imaged with both 50μm resolution T2*-weighted structural MRI and 1.2μm resolution μCT datasets centered on a coronal slice of brainstem and cerebellum. Using vasculature and neuronal somas as fiducials to guide the subsequent EM imaging (yellow arrows, Fig. 3d-e), we found individual synapses (white arrows, Fig. 3f) on neuronal somas in the EM data. Thus we traversed from MRI data of the entire coronal cerebellum/brainstem to underlying cellular architecture of the spinal vestibular nucleus with micron-resolution μCT data and ultra-structural details of individual synapses, all from the same brain.

**FIGURE 3.**
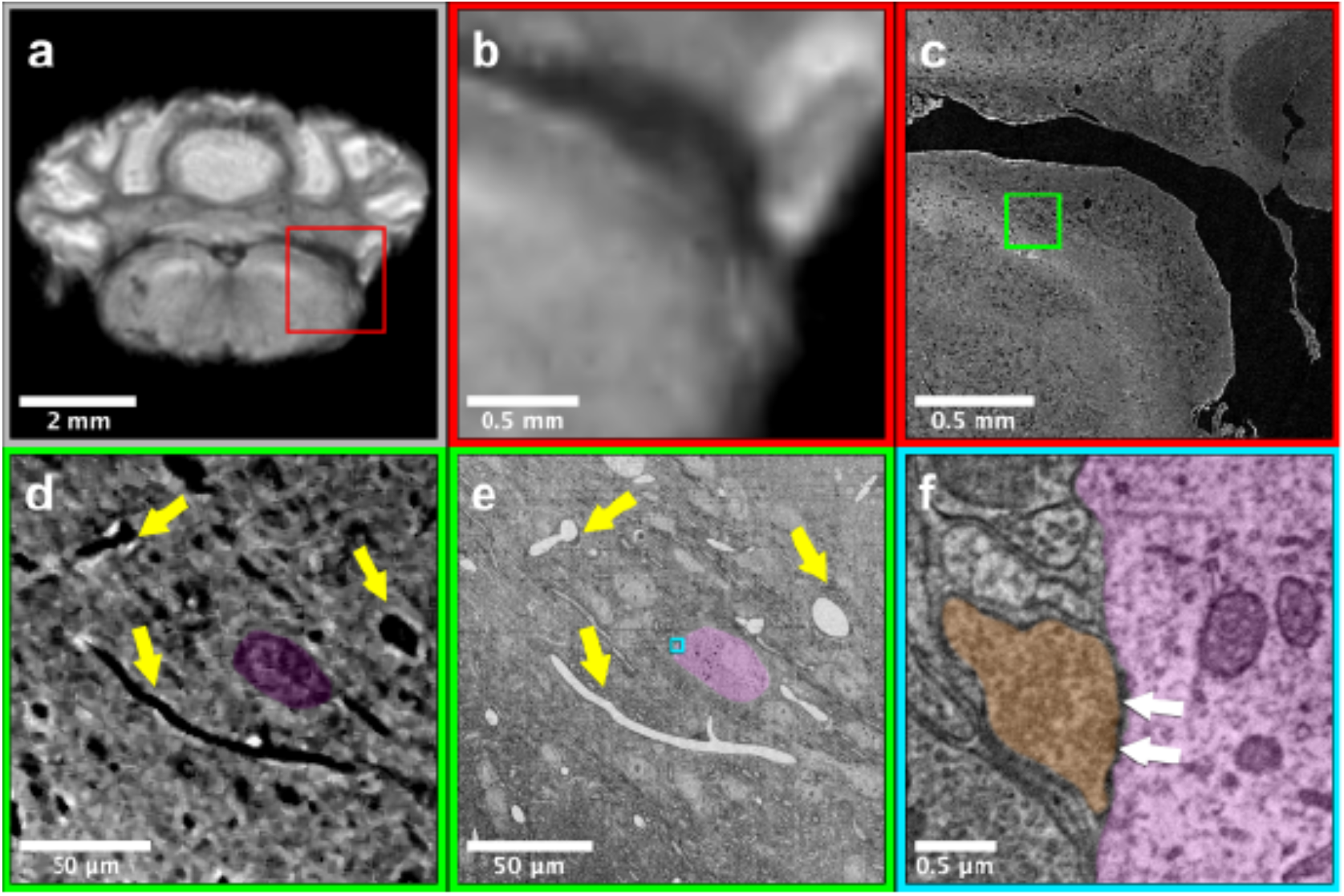
A multi-scale, multimodal pipeline for imaging the same brain from MRI to EM. The same brain was imaged using diffusion (a, b) MRI (50 micron isotropic voxels), (c, d) μCT (~ 1.2 um/voxel resolution), and (e, f) large volume serial electron microscopy (EM, ~ 3 nm/voxel resolution). (b, c) shows the same FOV from the whole brain MRI and μCT imaging, corresponding to the red ROI in (a). (d - e) show a smaller FOV in both the μCT and EM data, corresponding to the green ROI in (c). Yellow arrows indicate corresponding blood vessels, and a single neuron is labelled purple. Panel (f) highlights an individual somatic synapse (white arrows) on that soma, colored orange. The FOV of (f) is indicated by the blue ROI in (e). This pipeline demonstrates the ability to identify corresponding structures in a single brain imaged across four orders of magnitude of spatial resolution.

### 3.3 Comparing results of μCT with MRI

We next explored how whole brain μCT could indicate underlying cellular correlates of MRI signals. Since both approaches produce isotropic 3D renderings of whole volumes (Figs. 2a and 4b), this allowed for simple co-registration between the two datasets. For example, Figure 4a shows 3D somas/dendrites traced from x-ray data from the medial vestibular nucleus. While individually smaller than an MRI voxel, resolving the cytoarchitecture suggests a potential source of underlying contrast variability within an MRI dataset. Similarly, the somas in the dentate gyrus (DG) of the hippocampus are easily seen in the μCT data (Fig. 4c). Ten thousand individual cells were counted and traced within the DG over a 50μm slab of data. Again, while the individual somas are much smaller than the resolution of the MRI data, the high density of somas contribute to an anatomically dependent local variation in the MRI signal, rendering it resolvable from the surrounding structures (yellow box, Fig. 4b).

**FIGURE 4.**
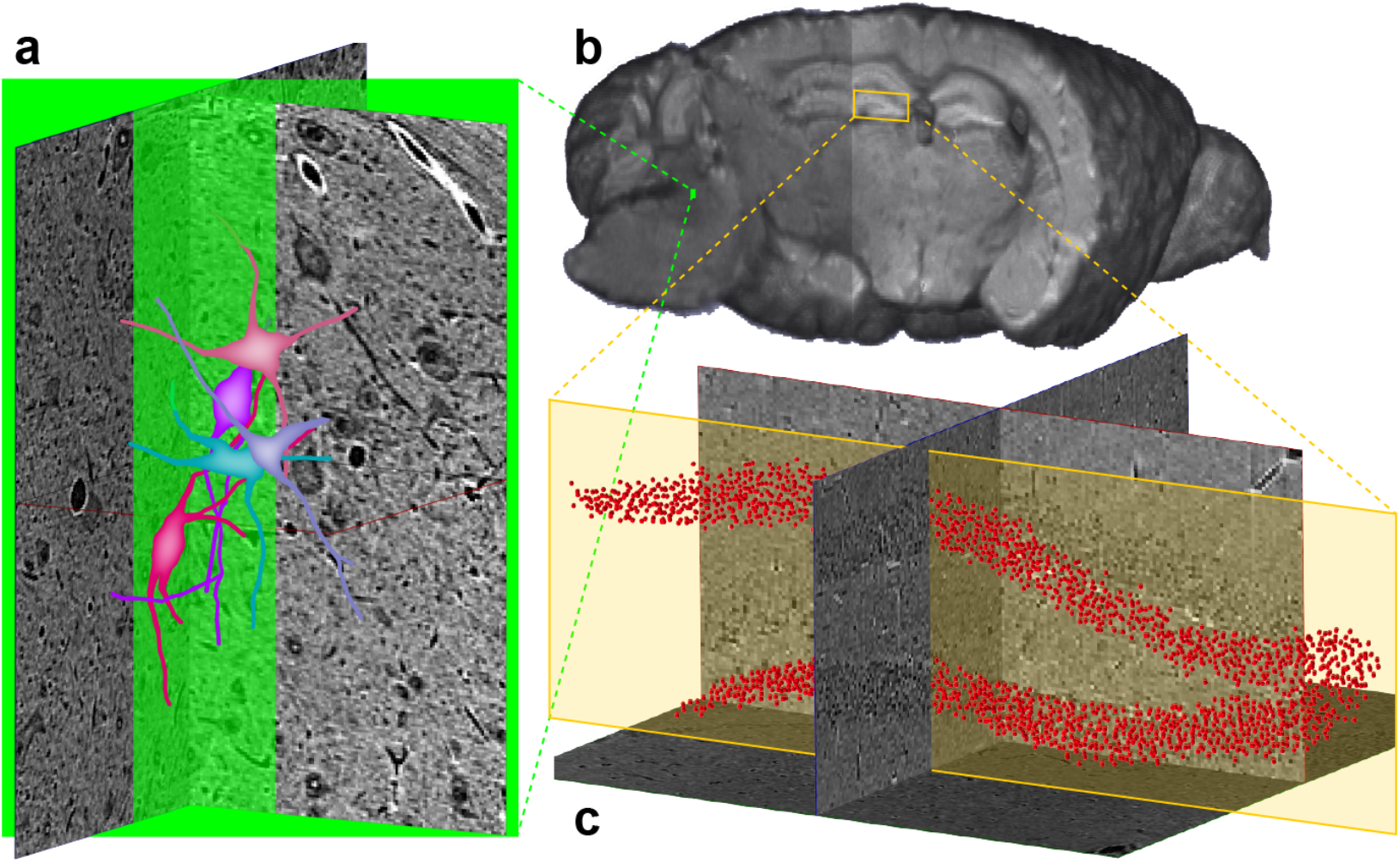
Whole brain synchrotron X-ray (μCT) provides cellular correlates of MRI contrast. (a) 3D rendering of structural MRI data used for coregistration with similar μCT data (a). Somas and dendrites of neurons in the medial vestibular nucleus (e) and somas in (g) dentate gyrus were traced from the μCT data. While these somas are much smaller than the resolution of the MRI data, they provide possible underlying sources of contrast in the T2* weighted images

Finally, we investigated DTI, which is explicitly sensitive to underlying axon distributions. We first segmented large myelinated tracts and vasculature in the μCT dataset using machine vision algorithms as a first pass comparison between μCT and DTI datasets and for further fine scale analyses. For example, Figures 4a and b show linearly co-registered sagittal cross-sections of the mouse cerebellum and brain stem in both structural MRI and μCT, respectively. Overlain on the MRI structural scan are estimates of crossing fibers in each voxel. Specifically, the MRI data supported estimates of two fiber populations (Fig. 5b): one population traversing an anterior/posterior (A/P) direction and another traversing the left/right (L/R) direction (green and red cylinders, respectively).

**FIGURE 5.**
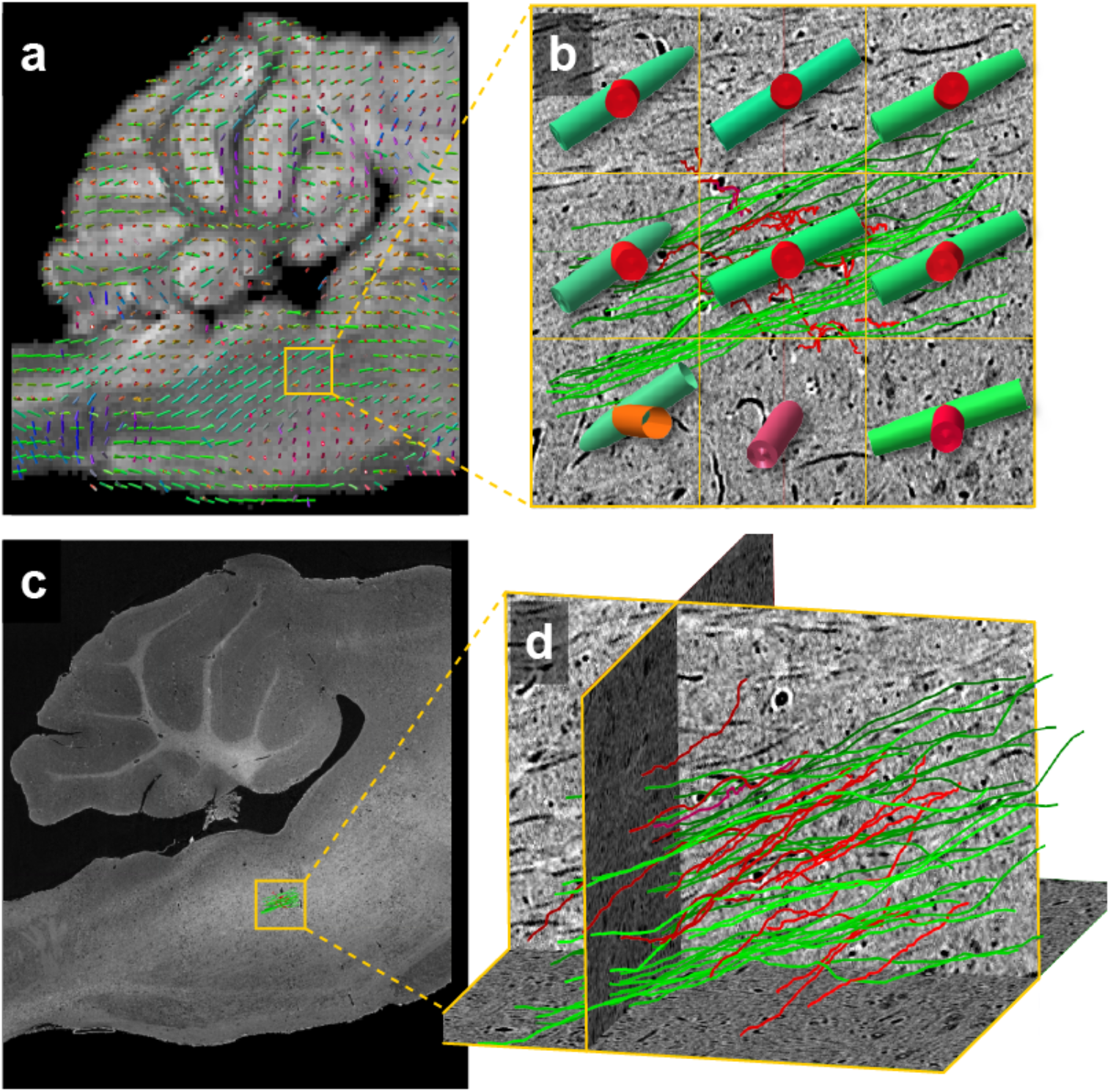
Whole brain synchrotron X-ray (μCT) provides axonal correlates of MRI contrast) (a) DTI data of the same brain were also acquired. Principal and secondary fiber population estimates are shown overlaid on a sagittal slice of a corresponding structural MRI scan (green indicates fiber orientation along the rostral/caudal axis, red indicates left/right) as well as in (b) the magnified 9 voxels. (c) The μCT data from the same brain has been linearly registered to the MRI structural data and (d) individually traced axons from the mCT data are shown with the same color scheme as (a) and (b). Both principal and secondary fiber estimates from DTI data indicate quite similar orientations as the underlying population of individual axons traced from the μCT data, as seen by (b) the agreement of the DTI direction estimates superimposed on the individual traced individual axons.

The individually traced axons from μCT were similarly color coded for comparison (Fig. 5d). The yellow boxes in Figures 5a and c indicate the location of 9 MRI voxels in which 47 individual axons (27 L/R, 20 A/P) were traced in the x-ray data. Figure 5a shows the MRI diffusion direction estimates overlain on top of the axons. The populations of axons traced in the x-ray data clearly follow similar directions to the MRI diffusion direction estimates while negligible numbers of axons in the x-ray data traverse directions other than two estimated with the DTI analyses. This supports the accuracy of the MRI diffusion model used to estimate directions of underlying fiber populations.

## 4. DISCUSSION

We describe a pipeline using three imaging modalities, providing unprecedented views across orders of magnitude of a single mouse brain sample. This will allow for potentially novel understandings and/or interpretations of results within each modality, currently constrained by scale-dependent differences in each individual approach. For example, for axon bundles, MRI data will provide low-resolution, whole-brain estimates of neural tracts (Fig. 1a), μCT will provide long range trajectories of individual myelinated axons in those tracts (Fig. 1b), validating and extending the MRI data, and EM would provide reconstructions of individual wrappings of myelin sheaths (Fig. 1c) or the eventual synaptic targets of those myelinated axons.

Because of the contrast mechanisms of each imaging modality, this pipeline works on brains (or any other organ type) of any species. MRI contrast is explicitly sensitive to tissue water, regardless of tissue type. Further, osmium is used as a contrast stain for both μCT and EM because it stains membranes of any type of cell in any type of tissue (Palay et al., 1962). Therefore, in a single brain sample, our pipeline reveals the relative locations of nearly every soma, blood vessel, myelinated axon and synapse. Other ultra-high resolution whole brain imaging approaches often rely on genetic expression of fluorescent proteins to label a subset (often < 1%) of neurons, limiting these approaches to a small range of tractable animals (i.e. mouse, fly, and worm (Chung and Deisseroth, 2013b)). Subset labeling, by necessity, precludes detailing how different brain structures, like neuronal somas, vasculature, or myelinated tracts, relate to each other. Finally, whole brain optical approaches, whether based on expansion (Chen et al., 2015) or optical index matching (Chung and Deisseroth, 2013a), often remove cell membranes, making them essentially incompatible with subsequent large volume electron microscopy.

Other clear advantages of using whole brain μCT as an intermediate approach is that datasets are: (1) isotropic resolution, facilitating unbiased tracing in any direction, (2) imaged through intact brains with minimal deformation, obviating computationally difficult large scale alignments (Chung and Deisseroth, 2013b; Saalfeld et al., 2012), and (3) compatible with sample preparation for downstream electron microscopy.

Such pipelines will likely be useful in studying brain pathology. Pathological alterations in neural function and behavior are well documented, but our understanding of their cellular underpinnings remains fragmentary. The complexity of brain circuitry and the marked heterogeneity of neuronal sizes and shapes across brains could mask even dramatic changes in specific neuronal subsets, making structural analyses difficult. Using μCT and EM to inform contrast variability in MRI data over entire brains would significantly narrow the search for pathological structural change and lead improved specificity in MRI. This would also lead to potential *in vivo* MRI biomarkers of disease. The next generation of synchrotrons currently under construction (Schmidt et al., 2018) should allow for even deeper penetration of samples with the possibility of imaging even larger brains in their entirety. Finally, by instantiating our imaging pipeline in a national laboratory, we provide a path for access for future investigations by other researchers at a no-cost facility open to researchers worldwide.

## 5. ACKNOWLEDGEMENTS

The authors thank Shawn Mikula for help with sample preparation for μCT and EM. S.F would like to thank the MRIS and MRIRC. N.K. and V.S. are supported from a Technical Innovation Award from the McKnight foundation, a Brain Initiative of the National Institutes of Health (U01 MH109100), and National Science Foundation Neuro Nex grant. P.LR. is partially supported by the National Institutes of Health (R01EB026300). S.T. is supported by the National Institute of Neurological Disorders and Stroke of the National Institutes of Health (F31NS113571). Additional funding supporting this work was provided by the National Institutes of Health (S10-OD025081, S10-RR021039, and P30-CA14599). This research used resources of the Advanced Photon Source, a U.S. Department of Energy (DOE) Office of Science User Facility operated for the DOE Office of Science by Argonne National Laboratory under Contract No. DE-AC02-06CH11357. This research also used resources of the Argonne Leadership Computing Facility, which is a DOE Office of Science User Facility supported under Contract DE-AC02-06CH11357. Finally, the authors would like to thank Viren Jain and Jennifer Neely for thoughtful comments and discussions over the course of the preparation of the manuscript.

## 6. ABBREVIATIONS

MRI: Magnetic Resonance Imaging
DTI: Diffusion tensor Imaging
EM: Automated Electron Microscopy
μCT: Synchrotron-based x-ray tomography

## 7. AUTHOR CONTRIBUTIONS

S.F., P.LR., and N.K. conceived and designed the imaging pipeline. S.F. acquired and processed MRI data. V.DA. designed the μ-CT system used for data collection and contributed in reconstruction of μ-CT data. V.S. collected all EM data. S.T. registered MRI, μCT, and EM data. S.F., A.S., and K.N. traced/segmented μCT data and S.F. registered results to MRI data. S.F. and N.K. wrote the final manuscript and all authors contributed to its revision.

