## Supplemental Information for "Multi-modal imaging of a single mouse brain over five orders of magnitude of resolution"

Supplemental Movie 1: **A downsampled whole mouse brain imaged at 1μm isotropic resolution with μCT**. Shown is a downsampled movie of the whole brain μCT dataset as described in the text. The isotropic data is shown from multiple perspectives and even downsampled, myelinated tracts (white bundles) and individual brain regions (e.g. cortex, hippocampus, etc.) can be clearly viewed. The ~10 terabyte dataset was collected in approximately 8 hours.

Supplemental Figure 1

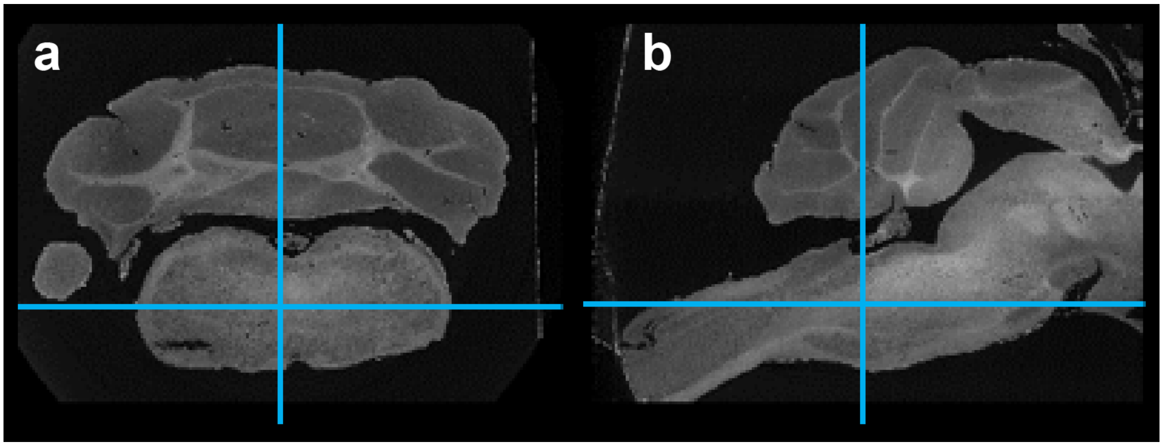


**The position in the medulla of supplemental movie 2.** Blue crosshairs in (a) coronal and (b) sagittal cross sections of the mouse brain indicate the region that supplemental movie 2 was taken from.

Supplemental Movie 2: **A sub-volume from the medulla of the whole brain imaged at 1μm isotropic resolution with μCT**. Shown is movie of a sub-volume of whole brain μCT dataset closer to full resolution, ~ 4x downsampled in plane (1.5mm x 1.5mm x 500μm). The data comes from the medial nuclei of the medulla. Individual neuronal somas (and nucleus and nucleolus) are clearly visible along and individual myelinated axons (small white tubes with grey rims) and blood vessels (large white tubes with dark black rims) can be traced long distances.
